# A cascade of structural rearrangements positions peptide release factor II for polypeptide hydrolysis on the ribosome

**DOI:** 10.1101/2025.03.09.642146

**Authors:** Nina Michael, Bridget Y. Huang, Korak Kumar Ray, Colin D. Kinz-Thompson, Ruben L. Gonzalez

## Abstract

Peptide release factor II (RF2) catalyzes the release of the nascent polypeptide from the bacterial ribosomal complex during translation termination and a subset of ribosome rescue pathways. Despite its critical role, the mechanisms that govern RF2 function and regulation remain elusive. Here, using single-molecule fluorescence energy transfer (smFRET), we characterize the conformational landscape that RF2 explores on the ribosomal complex and show that RF2 binding and dissociation from the ribosome follows a series of conformational rearrangements which depend on its ribosomal binding platform. We also show how further interactions with the ribosomal complex are necessary to properly position RF2 for polypeptide release. This work investigates not only the dynamics RF2 undergoes while in complex with the ribosome, but also identifies a potential mechanism by which the regulation of these dynamics may be disrupted, which may be exploited for future development of broad-spectrum antibiotics.

## MAIN TEXT

An essential step in the translation of messenger RNA (mRNA) into the corresponding encoded protein^1,2^ is the release of the newly synthesized polypeptide from the transfer RNA (tRNA) bound to the ribosomal peptidyl-tRNA binding (P) site. Polypeptide release is critical for the proper termination of the current round of translation and the efficient recycling of the ribosome for subsequent rounds. It thus plays an important role in both canonical translation termination at standard three-nucleotide ‘stop’ codons, and also in the rescue of ribosomes stalled on aberrant mRNAs, such as those lacking stop codons (‘non-stop’ transcripts).^3,4^

During canonical termination, a specific family of proteins called class I peptide release factors (RFs) recognize stop codons in the aminoacyl-tRNA binding (A) site in a termination complex (TC), where they bind and position a specific RF domain, domain III (dIII), into the ribosomal peptidyl transfer center (PTC) to facilitate the hydrolysis and release of the polypeptide chain.^5,6^ The ribosomal A site bearing the stop codon thus forms a binding platform that allows RFs to stably associate with TCs to perform their necessary function. This ability of RFs to recognize and stably bind to this ribosomal binding platform and subsequently undergo structural rearrangements to facilitate polypeptide release are critical parts of the mechanism of translation in all domains of life.^7^

In addition to its role in canonical translation termination, one of the two bacterial class I RFs, RF2, has been co-opted in certain bacterial species for the rescue of ribosomal complexes stalled on non-stop mRNAs.^8^ In the absence of a stop codon, the ribosome continues incorporating amino acids until the 3’ end of the non-stop mRNA enters the P site. This forms a non-stop elongation complex (nsEC) bearing an empty, or partially empty, A site.^9^ The nsEC lacks the necessary ribosomal binding platform capable of accommodating any aminoacyl-tRNA or RF and stalls on the mRNA. A secondary protein factor, alternative rescue factor A (ArfA), can recognize such complexes and form a substitute ribosomal binding platform for RF2 on the nsEC, facilitating peptide release and ribosome rescue.^10,11^ The ArfA-RF2 pathway serves as an important fail-safe for non-stop ribosome rescue and becomes essential for survival under conditions where other non-stop rescue pathways, such as trans-translation, are inactive.^4,12–14^ It has also been recently implicated as a necessary component in a newly-discovered bacterial phage defense pathway.^15^ Like canonical termination, the recognition and stable binding to ArfA-bound nsECs by RF2 and their subsequent structural transitions are critical steps for the efficient ArfA-mediated rescue and recycling of stalled nsECs. Furthermore, since the pathway of non-stop rescue is divergent between the different domains of life,^16^ the bacterial non-stop rescue pathways serve as attractive targets for the next generation of antibiotics.^17,18^

Multiple structural and biochemical studies have been employed that indirectly probe the mechanistic details of RF2 function in both canonical termination and ArfA-mediated non-stop rescue.^6,8,10,19–34^ These studies suggest that the individual steps of RF2-catalyzed peptide hydrolysis in these two distinct pathways are broadly similar. Both require a ribosomal binding platform for RF2 in the A site (comprising the stop codons UGA or UAA in TCs,^3^ and ArfA bound in the decoding center and the empty mRNA channel downstream of the A site in nsECs^11^). Upon stably associating with this binding platform, RF2 is expected to transition from a “collapsed” conformation, where dIII is far from the PTC, to an “extended” conformation, where dIII is “docked” into the PTC (Fig. 1).^19,20,22,23,26–29,31,32^ This structural transition is specifically thought to be triggered by specific RF2 interactions with the ribosome in TCs^19,20,23,32^ or ArfA in nsECs^26–29,31^ that stabilize an RF2 motif connecting domains II and III, termed the “switch loop”. An alanine to threonine mutation at position 18 in ArfA (mut-ArfA) leads to a loss of RF2 function,^12^ which structural studies have attributed to the disruption of switch-loop interactions believed to be essential for triggering the collapsed to extended transition in RF2.^27,28^

**Figure 1.**
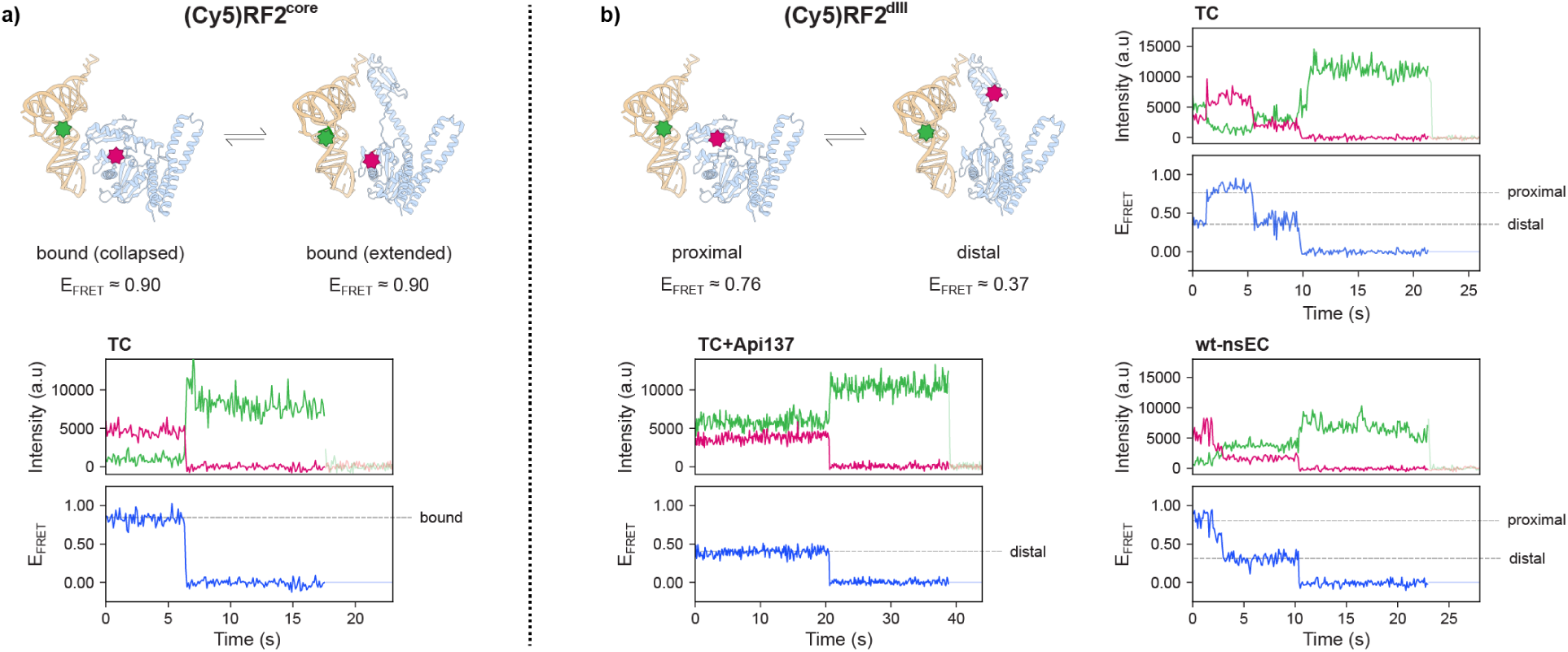
smFRET signals for RF2 stable association and dynamics. **(a)** <u>Top</u>: structural representations of RF2-bound ribosomal complexes containing the collapsed (PDB ID: 5U9G) and extended (PDB ID: 5U9F) conformations of RF2 with the relative labeling positions for Cy3 (green) and Cy5 (red) corresponding to the (Cy5)RF2^core^ smFRET signal and the expected E_FRET_ states. <u>Bottom</u>: a representative individual E_FRET_ trajectory for the TC showing the Cy3 (green) and Cy5 (red) fluorescence intensities and the corresponding E_FRET_ (blue). **(b)** <u>Top left</u>: structural representations of RF2-bound ribosomal complexes containing the collapsed and extended conformations of RF2 with the relative labeling positions for Cy3 (green) and Cy5 (red) corresponding to the (Cy5)RF2^dIII^ smFRET signal and the expected E_FRET_ states. <u>Top right and</u> <u>bottom</u>: representative individual E_FRET_ trajectories for the TC (top right), Api137-bound TC (bottom left), and wt-nsEC (bottom right), showing the Cy3 (green) and Cy5 (red) fluorescence intensities and the corresponding E_FRET_ (blue). For clarity, only RF2 and the P-site tRNA are displayed in all structures.

However, since the real-time conformational dynamics of RF2 during either canonical termination or ArfA-mediated non-stop rescue have not been directly observed, mechanistic models of RF2 function remain speculative. In this work, we use total internal fluorescence microscopy (TIRFM) coupled with single-molecule fluorescence resonance energy transfer (smFRET) to follow the real-time dynamics of the binding and conformational rearrangements of RF2 in ribosomal complexes. We develop and utilize two smFRET signals to investigate these RF2 dynamics in the context of TCs, nsECs bound to wild-type (wt-) ArfA (wt-nsECs), and nsECs bound to mut-ArfA (mut-nsECs). We demonstrate that ribosome-bound RF2 dynamically samples a wider conformational landscape than the previously defined collapsed and extended conformations. We show how these transiently sampled intermediate RF2 conformations are regulated by the ribosomal binding platform and, in mut-nsECs, form part of the mechanism through which mut-ArfA abrogates RF2 function. Finally, we elucidate a specific “docked” conformation of RF2 which must be stabilized by specific interactions with the ribosomal complex for peptide hydrolysis in both TCs and nsECs.

## Results

### Development of an smFRET signal to monitor RF2 binding to the ribosome

We first developed an smFRET signal to report on RF2 binding and dissociation from the ribosome. For the FRET donor, we used cyanine 3-labeled (Cy3-labeled) ribosomal complexes, labeled at a P site-bound phenylalanine-specific tRNA carrying a formylmethionine-phenylalanine dipeptide (fMet-Phe-(Cy3)tRNA^Phe^). For the FRET acceptor, we employed cyanine 5-labeled (Cy5-labeled) RF2, where RF2 domain II (hereafter referred to as the “core” domain) is labeled at residue 184 ((Cy5)RF2^core^). Since the core domain does not undergo significant structural rearrangements, the FRET efficiency (E_FRET_; defined as the Cy5 fluorescence intensity divided by the total Cy3 and Cy5 fluorescence intensities) for this signal is expected to remain the same (0.95–0.97, based on a FRET radius (*R*_0_) of 55 Å for Cy3-Cy5^35^) for the different ribosome-bound RF2 conformations previously observed in structural studies^28^ (Fig. 1a). Using this signal, we investigated RF2 binding to (*i*) elongation complexes (ECs) with a codon for lysine (AAA) in the A site, (*ii*) TCs with an A-site stop codon (UAA), and (*iii*) nsECs containing a 3’-truncated mRNA with an empty A site, in the presence of both wt- and mut-ArfA (see Methods for details).

As expected, steady-state experiments of (Cy5)RF2^core^ binding to TCs and nsECs yielded one non-zero E_FRET_ state centered at ∼0.9 (Figs. 1a, S1), according to a hidden Markov model (HMM)-based analysis^36^ (see Methods for details). This non-zero E_FRET_ state is consistent with stable binding of RF2 to both TCs and nsECs. These experiments also yielded a zero E_FRET_ state centered at 0.0, which corresponds to either (Cy5)RF2^core^ being too far away from the P-site tRNA to yield any observable E_FRET_ (*e.g.*, when RF2 is not bound to the ribosome) or the Cy5 acceptor fluorophore being photobleached (Fig. 1a). When performing similar steady-state experiments of (Cy5)RF2^core^ binding to ECs, we only observed the zero E_FRET_ state (data not shown). Since RF2 must interact with the ribosomal complexes to probe the identity of the A site binding platform and distinguish between ECs, TCs, and nsECs, these results suggest that a transient “encounter complex”^37^ is formed when RF2 first interacts with the ribosome, and that it has a lifetime much shorter than our TIRFM acquisition time of 100 ms (see Methods). For TCs and nsECs, the presence of the ribosomal binding platforms stabilize the encounter complex and leads to more stably bound and long-lived RF2 conformations that correspond to the observed non-zero E_FRET_ state. For ECs, the lack of such a platform in the A site means RF2 cannot stably bind, and so it rapidly dissociates.

### TCs facilitate the stable association of RF2 more effectively than wt-nsECs and mut-nsECs

We next employed the core smFRET signal in pre-steady-state experiments where we delivered (Cy5)RF2^core^ to TCs and wt-nsECs. For wt-nsECs, a saturating concentration of wt-ArfA (more than 50-fold excess of the (Cy5)RF2^core^ concentrations) was added along with (Cy5)RF2^core^ to the ribosomal complexes immediately prior to imaging (see Methods for details). By titrating the concentration of (Cy5)RF2^core^ over the course of several successive pre-steady state experiments, we were able to obtain the second-order rates of association (*k*_*a*_s) of RF2 to TCs and wt-nsECs (Fig. S2a).

The *k*_*a*_ of (Cy5)RF2^core^ for TCs was 12.6 ± 0.7 (μM·s)^−1^. In the absence of wt-ArfA, we did not observe any stable (Cy5)RF2^core^ binding to nsECs (data not shown); however, in the presence of wt-ArfA, the *k*_*a*_ of (Cy5)RF2^core^ for the wt-nsECs was 7.1 ± 0.7 (μM·s)^−1^ (Fig. S2b). Because the rate of formation of the initial encounter complex between (Cy5)RF2^core^ and the ribosome depends on the collision rate between the two, which should be approximately the same for both TCs and nsECs,^37^ the *k*_*a*_s that we measured instead reflect the probability that the encounter complex progresses to a stable and long-lived RF2-bound ribosomal complex. The nearly two-fold larger *k*_*a*_ of the TC over that of the wt-nsEC thus suggests that the TC binding platform facilitates productive association of RF2 more successfully than the wt-nsEC binding platform.

For mut-nsECs, we estimated the *k*_*a*_ from steady-state experiments as 7.2 ± 0.3 (μM·s)^−1^ (Fig. S2b). Therefore, mut-ArfA does not appear to reduce RF2 stable association with ribosomal complexes when compared to wt-ArfA, which is in agreement with previous studies.^10,12^ The ability of mut-ArfA to abrogate RF2 function must therefore arise from the dysregulation of a mechanistic step that occurs after RF2 association with the mut-ArfA-bound ribosomal binding platform.

### ArfA must accommodate into the A site to form a functional ribosomal binding platform for RF2

We next investigated the role of ArfA in forming a stable ribosomal binding platform for RF2 on nsECs. Previous studies have hypothesized that ArfA must adopt a specific conformation on nsECs before RF2 can stably associate.^8,11^ We therefore performed steady-state smFRET experiments where we allowed wt-ArfA to accommodate into the nsEC A site before the addition of (Cy5)RF2^core^. When a saturating concentration of wt-ArfA was added to nsECs and pre-incubated for 5 min prior to the addition of (Cy5)RF2^core^, the rate of RF2 dissociation (*k*_*d*_) from the resulting wt-pre-nsECs (where ‘pre’ denotes the pre-incubation) was 0.26 ± 0.03 s^−1^ (Table 1). In contrast, when a saturating concentration of wt-ArfA was added simultaneously with (Cy5)RF2^core^, the *k*_*d*_ of RF2 from the resulting wt-nsEC was 0.66 ± 0.05 s^−1^ (Table 1), demonstrating that RF2 is less stably bound to wt-nsECs than wt-pre-nsECs. This supports the hypothesis that ArfA accommodates in the A site first prior to RF2 stable association.

**Table 1.** Dissociation rates and conformational rearrange rates for ribosome-bound RF2. The results of the HMM-based analysis for the dissociation rate (*k*_*d*_) of (Cy5)RF2^core^ for different ribosomal complexes, and for the rates of the proximal-to-distal transition (*k*_*p*→*d*_), the distal-to-proximal transition (*k*_*d*→*p*_), the dissociation from the proximal state 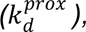 and the dissociation from the extended state 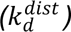 of (Cy5)RF2^dIII^ for different ribosomal complexes. All dissociation rates except for 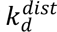 of (Cy5)RF2^dIII^ from the Api137-bound TC (indicated by *) are corrected for photobleaching. Errors represent the standard deviation of the posteriors inferred from the HMM-based analysis.

|  | (Cy5)RF2<sup>core</sup> | (Cy5)RF2<sup>dIII</sup> |  |  |  |
| --- | --- | --- | --- | --- | --- |
| | $k_d$ (s<sup>-1</sup>) | $k_{p \rightarrow d}$ (s<sup>-1</sup>) | $k_{d \rightarrow p}$ (s<sup>-1</sup>) | $k_d^{prox}$ (s<sup>-1</sup>) | $k_d^{dist}$ (s<sup>-1</sup>) |
| **TC** | 0.16 ± 0.02 | 0.50 ± 0.02 | 0.35 ± 0.01 | 0.13 ± 0.02 | 0.41 ± 0.01 |
| **wt-nsEC** | 0.66 ± 0.05 | 0.35 ± 0.02 | 0.32 ± 0.01 | 0.08 ± 0.02 | 0.30 ± 0.02 |
| **wt-pre-nsEC** | 0.26 ± 0.03 | 0.32 ± 0.02 | 0.25 ± 0.01 | 0.08 ± 0.02 | 0.18 ± 0.01 |
| **mut-nsEC** | 0.47 ± 0.03 | 0.40 ± 0.02 | 0.33 ± 0.02 | 0.00 ± 0.01 | 1.11 ± 0.03 |
| **Api137-bound TC** | - | - | - | - | 0.24 ± 0.01* |

The *k*_*d*_ of RF2 from mut-nsECs was 0.47 ± 0.03 s^-1^, a value that is lower than that from wt-nsECs but nearly double that from wt-pre-nsECs (Table 1). The similar *k*_*d*_s of RF2 dissociating from mut-nsECs and wt-nsECs provides further evidence that mut-ArfA can also form a ribosomal binding platform for RF2 in mut-nsEC.

In the case of TCs, we observed an RF2 *k*_*d*_ of 0.16 ± 0.02 s^−1^, a rate that is lower than the RF2 *k*_*d*_ for any nsEC (Table 1). This indicates that the TC forms a ribosomal binding platform that is much more effective at binding RF2 than the platform formed by wt-ArfA, even for the wt-pre-nsEC, at least under our experimental conditions. Notably, the *k*_*d*_ of RF2 from TCs that we observe is almost an order of magnitude lower than that reported in a previous study,^21^ suggesting that RF2 binds stably enough to TCs such that a secondary factor (*e.g.*, RF3) would be required for rapid RF2 dissociation (see Discussion).

### Development of an smFRET signal to follow RF2 conformational dynamics

We next investigated the conformational dynamics of the ribosome-bound RF2 using a separate smFRET signal. This signal used a different RF2 construct as a FRET acceptor. Here, RF2 was labeled with Cy5 at residue 271 within dIII, the domain that extends into the PTC to carry out peptide hydrolysis ((Cy5)RF2^dIII^) (Fig. 1b). We expected this smFRET signal to report on RF2 structural rearrangements that resulted in the repositioning of dIII, such as the ones captured in previous structural studies^28^ (Fig. 1b).

We observed two non-zero E_FRET_ states for (Cy5)RF2^dIII^ for both TCs and nsECs (Fig. 1b, S1b): a low-E_FRET_ state of ∼0.37 for both TCs and nsECs, where RF2 dIII is distal to the Cy3-labelled tRNA, and a high-E_FRET_ state of 0.76 for TCs and 0.70 for nsECs, where RF2 dIII is proximal to the Cy3-labelled tRNA (Table S1). We expected these “distal” and “proximal” E_FRET_ states to respectively correspond to the extended and collapsed conformations previously observed in structural studies^28^ (Fig. 1b). However, the observed E_FRET_ states were significantly different from what we expected from the above conformations (0.64 and 0.96 for the extended and collapsed conformations, respectively, using a Cy3-Cy5 R_0_ value of 55 Å). While such a difference might be due to changes in the photophysical properties of the fluorophores in the ribosomal environment, it was not readily obvious whether or how the conformations of RF2 that gave rise to the observed E_FRET_ states in our smFRET experiments corresponded to those captured in the structural studies. Since our smFRET signal reports only on the distance between the FRET donor and acceptor, we adopted the labels “distal” or “proximal” (relative to the Cy3 label on the P-site tRNA, Fig. 1b) for our E_FRET_ states to avoid assigning structural interpretations until a more detailed analysis could be carried out.

To assign these observed E_FRET_ states, we used an antimicrobial peptide, Api137, which binds the nascent polypeptide exit tunnel of a TC post-peptide-hydrolysis and traps RFs in an extended conformation.^38,39^ In our experiments, TCs in the presence of Api137 primarily adopted a single non-zero low-E_FRET_ state centered at 0.39 (Fig. 1b), which we therefore assigned to the above extended conformation in the Api137-bound TC. We reasoned that this RF2 conformation also yielded the similar distal low-E_FRET_ state (centered at 0.37) in the non-Api137 bound TCs, wt-nsECs, and wt-pre-nsECs. A low-E_FRET_ state was also observed in the mut-nsECs. However, this was centered at 0.43, suggesting that this state was distinct from the distal states observed for TCs and wt-nsECs. This indicates the existence of at least two ribosome-bound RF2 conformations that can yield similar distal E_FRET_ states in our smFRET experiments, thus requiring further characterization.

The proximal E_FRET_ state from the other complexes was virtually absent for the Api137-bound TC (Fig. 2). Since this complex is expected to not adopt a long-lived collapsed RF2 conformation, we could therefore assign the proximal state to this collapsed conformation. The slight differences in E_FRET_ for this state in a wt-nsEC (0.70) compared to that in a TC (0.76) or a mut-nsEC (0.75) suggest subtle variations in the positioning of RF2 dIII between ribosomal complexes. However, similar to the distal E_FRET_ state, further characterization of the proximal E_FRET_ state is required to determine if it corresponds to a single or multiple RF2 conformations.

**Figure 2.**
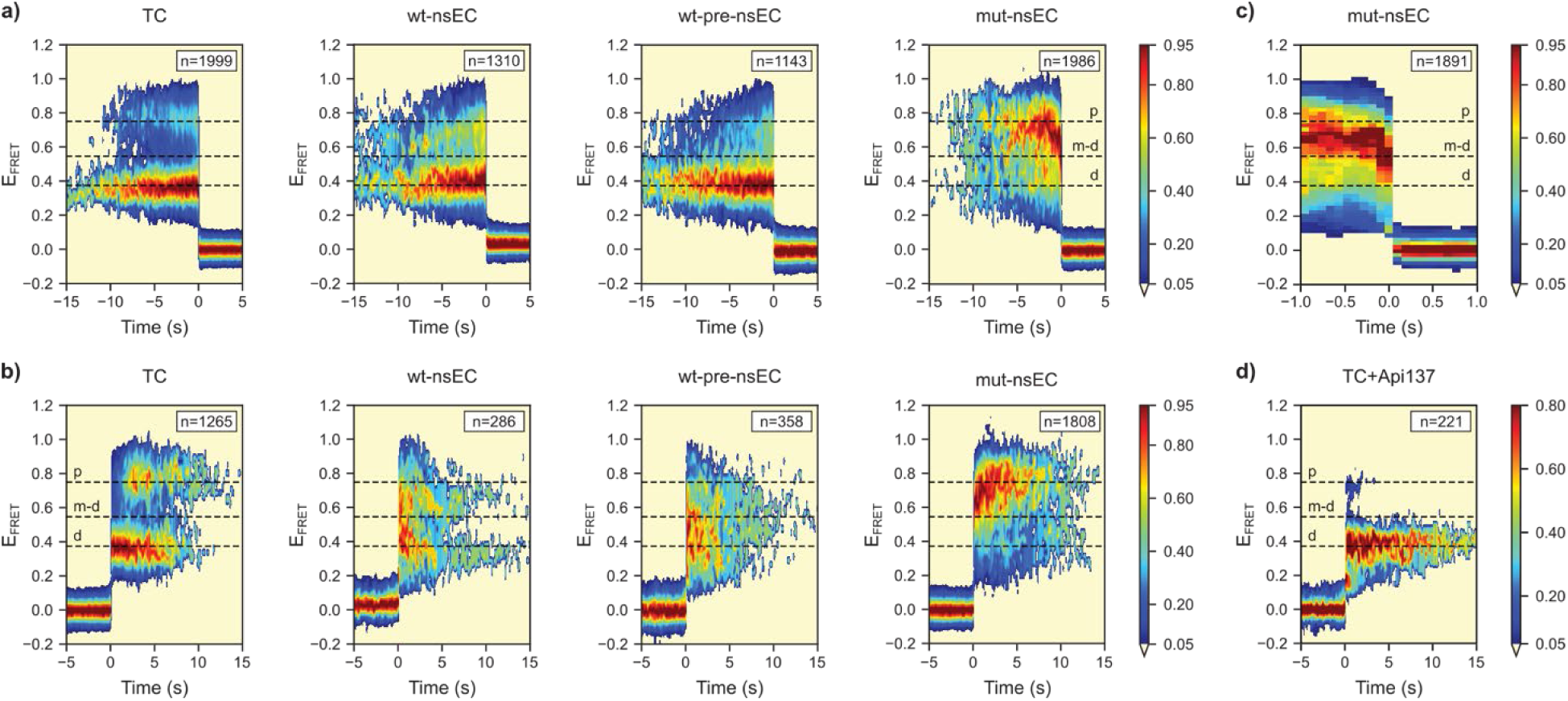
2D E_FRET_ vs time histograms of RF2 dissociation and stable association. 2D histograms post-synchronized to **(a)** the transition from any non-zero E_FRET_ state to the zero E_FRET_ state, following the dissociation of ribosome-bound (Cy5)RF2^dIII^ from TCs, wt-nsECs, wt-pre-nsECs, and mut-nsECs, **(b)** the transition from the zero E_FRET_ state to any non-zero E_FRET_ state, following the stable association of (Cy5)RF2^dIII^ to TCs, wt-nsECs, wt-pre-nsECs, and mut-nsECs, **(c)** the transition from any non-zero E_FRET_ state to the zero E_FRET_ state for mut-nsECs which highlights the transient mid-distal state adopted immediately prior to RF2 dissociation, and **(d)** the transition from the zero E_FRET_ state to any non-zero E_FRET_ state, following the stable association of (Cy5)RF2^dIII^ to Api-137-bound TCs, highlighting the excursion to the proximal E_FRET_ state in these complexes. Dashed lines show the E_FRET_ states corresponding to proximal (p), distal (d), and mid-distal (m-d) conformations of (Cy5)RF2^dIII^.

Since Api137 traps RFs in the distal state for time intervals much greater than our experimental time window,^38,39^ all instances of a transition from this state to the zero E_FRET_ state in the Api137-bound TC were likely due to photobleaching of (Cy5)RF2^dIII^. The HMM-derived rate of transition from this E_FRET_ state to the zero E_FRET_ state thus provided a conservative estimate of the rate of Cy5 photobleaching in our experimental conditions. All experimental *k*_*d*_s reported in this study were therefore corrected for photobleaching using this estimated photobleaching rate (see Methods for details).

### RF2 adopts at least two P-site tRNA distal conformations when bound to ribosomal complexes

The differences in the E_FRET_ states for different ribosomal complexes suggests that the ribosome-bound RF2 explores a wider ensemble of conformations than what previous studies have reported. In fact, we observed RF2 structural rearrangements in all four ribosomal complexes that we studied (Fig. 2). This was in striking contrast to a previous structural study of mut-ArfA which had suggested that the mutation in ArfA inhibits RF2 structural transitions and traps RF2 in a collapsed conformation.^27^ If RF2 was trapped in a collapsed conformation in mut-nsECs, we would only observe the proximal state in our experiments. Furthermore, our HMM-derived rates for transitions between the distal and proximal states were very similar for all complexes (Table 1). Thus, the rates of RF2 structural rearrangements were, on average, unaffected by the identity of the specific ribosomal complex. This intrinsically dynamic behavior of the ribosome-bound RF2 appears to agree with previous studies hypothesizing that RF2 dynamics may be regulated by changes in the local ribosomal environment that are triggered by RF2 binding.^30,40^ However, we have previously shown that HMMs do not distinguish between the distinct kinetic phases that arise from conformational transitions between states that have the same E_FRET_ values but that occur at different rates; instead, HMMs only report the average of these rates.^36^

To investigate the existence of any distinct kinetic phases, we instead quantified the dwell times distributions of the distal state in the E_FRET_ trajectories (*i.e.*, the times spent in the distal state before transitioning to the proximal state). Since the presence of distinct kinetic phases would manifest as non-single-exponential decays, we fit the resulting survival curves of the dwell time distributions to both single- and double-exponential decays (see Methods for details). To distinguish between the single- or double exponential fits, we used a cutoff value of 0.95 for the goodness-of-fit parameter, R^2^, for supporting the single-exponential decay. Using this criterion, the observed kinetics of the TC-bound RF2 could be adequately explained by a single exponential decay (R^2^ = 0.97) (Fig. S4a), but they needed at least a double-exponential decay to be properly modeled for nsECs (with R^2^ values of 0.94, 0.92, and 0.83 for the single-exponential fit for wt-nsECs, wt-pre-nsECs, and mut-nsECs, respectively) (Fig. S4a). Thus, our data show that the distal state encompasses at least two distinct RF2 conformations with indistinguishable E_FRET_ values, one of which corresponds to the extended conformation captured in structural studies. In the case of TCs, we note that it is possible that the corresponding RF2 conformations transition with similar rates, thereby giving rise to an apparent single-exponential decay.

Strikingly, when we constructed post-synchronized, two-dimensional (2D) E_FRET_ histograms of this distal state (where the transitions either to this state from the zero E_FRET_ state or the reverse is aligned to the zero time-point) (Fig. S3), we see the clear appearance of one such additional state for the mut-nsEC. This transiently sampled E_FRET_ state appears to be centered around 0.55, distinct from the distal states we previously observed. We attribute this state to a conformation of RF2 where dIII is more distal to the P-site tRNA than the collapsed conformation observed in structural studies. Hereafter, we refer to this E_FRET_ state as the mid-distal state. We reason that the transient nature of the corresponding RF2 conformation and the proximity of its E_FRET_ value to the low-E_FRET_ distal state prevented its identification as a separate state by our HMM-based analysis,^36^ which instead assigned it as a part of the distal state.

In contrast to the distal states, the corresponding observed kinetics for the proximal state for all complexes could be well explained by a single-exponential decay (with R^2^ greater than 0.95), suggesting the proximal state represents a single RF2 conformation (Figs. 3b and S5b). We reason that this proximal state must then arise from the collapsed RF2 conformation observed in structural studies. Unlike the distal states above, we can therefore use the terms “proximal” and “collapsed” interchangeably to refer to the same E_FRET_ state in our experiments. However, we have chosen, for consistency, to use proximal state for discussing our subsequent results.

**Figure 3.**
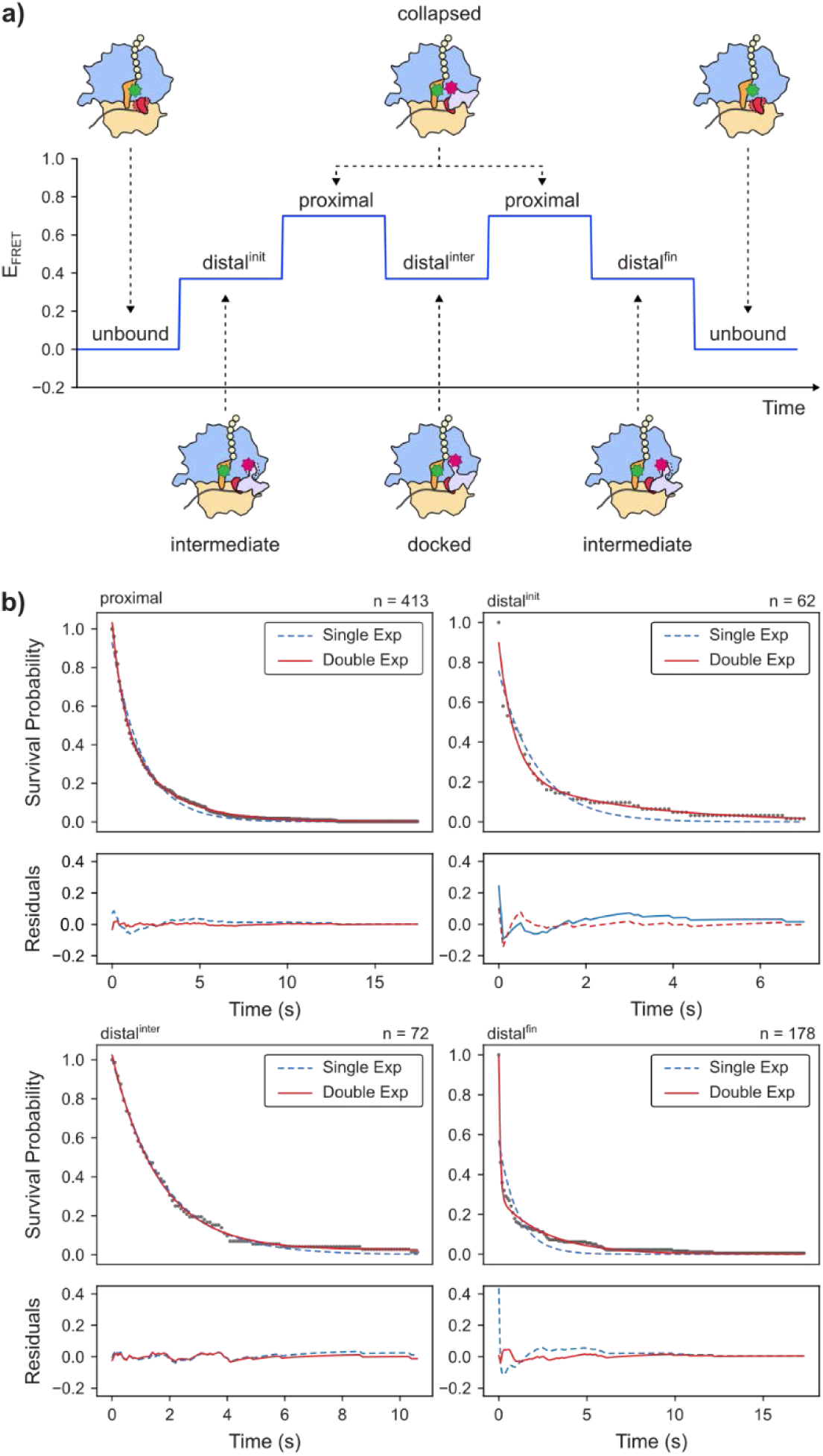
Dwell-time analysis of ribosome-bound (Cy5)RF2^dIII^. **(a)** A hypothetical E_FRET_ trajectory of (Cy5)RF2^dIII^, showing the different observed E_FRET_ states and the corresponding RF2 conformations. **(b)** Survival plots of the dwell time distributions (black) for wt-pre-nsECs along with fits of single-exponential (blue) and double-exponential (red) decays (above) and the corresponding residuals (below) for proximal, distal^init^, distal^inter^, and distal^fin^ states.

### RF2 dissociates from ribosomal complexes via previously undetected intermediate states

Having revealed the presence of at least two distal RF2 conformations, and in particular, their emergence when RF2 transitions to the zero E_FRET_ state, we next sought to investigate their mechanistic significance to RF2 function. We quantified the preference of RF2 to dissociate from either a distal or proximal state using steady-state smFRET experiments with (Cy5)RF2^dIII^ (Fig. 1b, S1b), which yielded the RF2 dissociation rate from these states (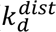 and 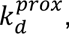 respectively) (Table 1). For all complexes that we investigated, 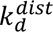 was significantly greater than 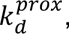 suggesting that RF2 preferentially dissociates from the ribosome in a distal state. Since the rates of exchange between the two states were at least an order-of-magnitude lower than our acquisition rate (Table 1), we could discount the possibility that the ribosome-bound RF2 transiently samples the proximal state immediately prior to dissociation for a period that does not show up in our experiments.^41^ This preference for RF2 to dissociate from the distal state was most dramatically pronounced in mut-nsECs, where 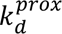 was negligible and 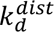 was 1.11 ± 0.03 s^−1^, which was more than 2-fold higher than that of TCs (0.41 ± 0.01 s^−1^) and 4- to 6-fold higher than that for wt-nsECs (0.30 ± 0.02 s^−1^ for wt-nsECs and 0.18 ± 0.01 s^−1^ for wt-pre-nsECs).

By synchronizing every transition from a non-zero E_FRET_ state to the zero state and visualizing those transitions using 2D E_FRET_ histograms (see Methods), we observed that RF2 preferentially dissociates from the mid-distal state in mut-nsECs (Fig. 2a). The mid-distal state exhibits a very short lifetime of <300 ms, which corresponds to a *k*_*d*_ > 3.3 s^−1^ (Fig. 2c). For nsECs, we saw an increased density of E_FRET_ values between the proximal and distal states, suggesting the presence of one or multiple transient mid-distal states (Fig. 2a, S3a). For these complexes, this mid-distal state is even more transiently sampled and, thus, even less likely to be identified by our HMM-based analysis.^36^ We surmise that it is this density that is simply significantly increased in mut-nsECs, leading to the observation of a distinct observed E_FRET_.

To more confidently detect these intermediate conformations in the dissociation pathway of RF2 from TCs and wt-nsECs, we employed the dwell-time based analysis described above. We specifically selected only those dwells in a distal E_FRET_ state that are preceded by the proximal state and followed by the zero E_FRET_ state. We expect these dwells to report on the final behavior of RF2 immediately before dissociation (distal^fin^) (Fig. 3a). For these distal^fin^ dwells, all complexes required at least a double-exponential fit (Fig. 3b, S5, and Table S2), suggesting that the mid-distal conformation and, likely, other distal conformations with closely spaced E_FRET_ values form an ensemble of transient intermediates that RF2 samples during dissociation from TCs and nsECs. This ensemble nature is supported by the broad distribution of E_FRET_ values observed immediately prior to RF2 dissociation from these complexes (Fig. 2a). Further studies are required to ascertain the structural details of these conformations.

### The binding intermediates of RF2 are regulated by the ribosomal binding platform

Having found that RF2 dissociates from the ribosomal complex through an ensemble of different intermediates, we next investigated if RF2 association to ribosomal complexes also occurs through a similar ensemble. Like above, we visualized the RF2 binding pathway using post-synchronized 2D E_FRET_ histograms where every transition from the zero state to any non-zero E_FRET_ states was aligned (Fig. 2b). Density corresponding to mid-distal state is present in the RF2 binding pathway for both mut-nsECs and wt-nsECs, but less so for wt-pre-nsECs. For TCs, on the other hand, this density is virtually absent; instead, we see that the distal state is immediately sampled upon stable RF2 association before the transition to the proximal state. This immediate sampling of the distal state upon stable RF2 association is also observed in the post-synchronized 2D E_FRET_ histogram for the Api137-bound TC complex (see above). For these complexes, sampling of the distal state is then followed by a transient excursion to the proximal state and a subsequent final excursion to a long-lived distal state, which represents the Api137-trapped extended state of RF2 after peptide hydrolysis (Fig. 2d). Taken together, this demonstrates that the pathway of RF2 association proceeds via a distal state for TCs and an ensemble of distal and mid-distal intermediate states for nsECs, which then transition to a proximal (collapsed) conformation before presumably transitioning to one or more hydrolysis competent distal conformations (see below, Fig. 3a).

Evidence for this binding pathway is also present in the dwell-time distributions of the distal states immediately following RF2 stable binding (distal^init^). These distal^init^ dwells are preceded by the zero E_FRET_ state and followed by the proximal state, and thus report on the initial behavior of the ribosome-bound RF2 (Fig. 3a). The distal^init^ dwell-time distribution was well-fit by a single exponential for TCs (R^2^ = 0.96), but not for nsECs (R^2^ values of 0.84, 0.90, and 0.89 for wt-nsEC, wt-pre-nsEC, and mut-nsEC respectively), which, similar to the distal^fin^ dwell-time distributions, required at least a double-exponential fit (Figs. 3b, S5, and Table S2). This suggests that RF2 also samples the ensemble of transient intermediates when binding to nsECs.

Altogether, our data suggests that the ribosomal binding platform regulates the ensemble of intermediate states sampled by RF2 during stable association. Additionally, while mut-ArfA can form a ribosomal binding platform, the specific nature of this binding platform makes RF2 more prone to populating the mid-distal state, which subsequently leads to its dissociation from the mut-nsEC. The ensemble of distal and mid-distal states thus form critical intermediates that are sampled during stable association and dissociation of RF2 (see Discussion).

### The hydrolysis-competent docked state of RF2 is distinct from the distal states that RF2 samples during association and dissociation

While our results demonstrate that an ensemble of distal intermediates regulates RF2 binding and dissociation, it was not immediately clear how these intermediates are related to the extended conformation observed in structural studies. We found that the distribution of distal^inter^ dwells (distal states preceded or followed by the proximal state) could be reasonably explained as a single-exponential for all ribosomal complexes except mut-nsEC (Figs. 3b, S5, and Table S2). Since these dwells exclusively report on the behavior of ribosome-bound RF2 distal conformations adopted after the proximal state (Fig. 3a), and are thus distinct from the above ensemble of binding intermediates, we propose that this one distal RF2 conformation corresponds to a docked conformation in which the RF2 dIII is extended into the PTC and competent for peptide hydrolysis. We hereafter refer to this as the “docked” conformation.

For mut-nsECs, the distal^inter^ dwell-time distribution was not well explained by a single-exponential decay (R^2^ = 0.94) (Fig. S5, Table S2); however we did not observe many dwells for this state (n=29) relative to the other ribosomal complexes. The distribution from the dwells we do observe suggests that the mut-nsECs distal^inter^ state more likely resembles the distal and mid-distal states observed during association and dissociation, although possibly with some transient sampling of the docked conformation. Thus, it is likely that the disruption of the ability of RF2 to properly dock into the PTC leads to the increased occupancy of the mid-distal conformation (and other binding intermediates) in mut-nsECs, which subsequently results in the rapid dissociation of RF2.

## DISCUSSION

The smFRET experiments we presented here have probed the entire mechanistic pathway of RF2 stable association and conformational dynamics leading up to RF2-catalyzed peptide hydrolysis for two distinct biological processes: canonical translation and ArfA-mediated non-stop rescue. Our results reveal that RF2 associates with the ribosome through an ensemble of binding intermediates that all lead to a collapsed conformation, and then to a hydrolysis-competent docked conformation (Fig. 4a and 4b). We showed that the initial transitions between the binding intermediates and the collapsed conformation are regulated by the ribosomal binding platform for RF2, and that they occur even in the presence of mut-ArfA. However, the docked state that appears following the second transition is likely stabilized by further interactions within the ribosomal complex, including those with the RF2 switch loop, as suggested in previous studies^19,20,23,26–29,31,32,42^. It is the disruption of these interactions in mut-nsECs that abrogates the ability of RF2 to remain stably docked into the PTCs in this complex.

**Figure 4.**
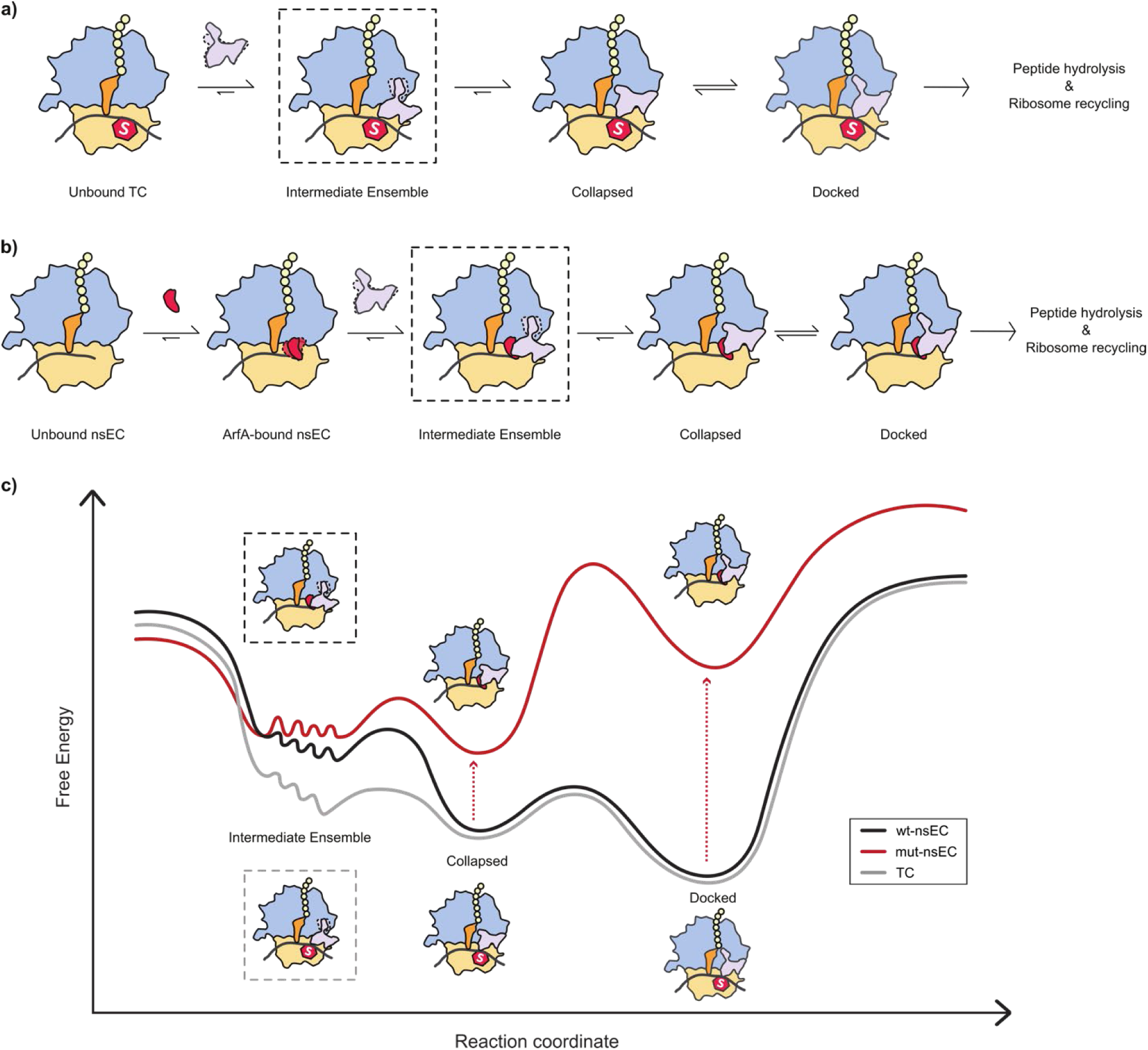
Proposed model for the mechanism of RF2 function. **(a)** RF2 associates with the TC in an ensemble of binding intermediates, subsequently transitions to the collapsed conformation, and then docks into the PTC, allowing it to hydrolyze the newly synthesized polypeptide. **(b)** ArfA stably associates with the nsEC, followed by RF2 associating in an ensemble of binding intermediates, subsequently transitioning to the collapsed conformation, and then docking into the PTC, allowing it to hydrolyze the aberrant, truncated polypeptide. **(c)** A schematic of the RF2 conformational free energy landscapes on wt-nsECs (black), mut-nsECs (red) and TCs (grey) along with the corresponding ribosome-bound RF2 conformations, showing the cascade of structural transitions that allow RF2 to achieve the docked conformation in wt-nsECs and TCs, and the perturbation caused by mut-ArfA to the landscape that reverses the cascade and abrogates RF2 function in mut-nsECs.

We hypothesize that upon initial formation of an RF2-ribosome encounter complex, RF2 cascades through multiple intermediate conformations, and that this ensemble of transiently populated RF2 conformations allows RF2 to efficiently undergo stochastic transitions to the more thermodynamically stable collapsed and docked conformations observed in structural studies (Fig. 4c). mut-ArfA disrupts this cascade by stabilizing the intermediate distal conformations relative to the collapsed conformation (Fig. 2), and thus inhibits transitions into the docked conformation for mut-nsECs (Figs. 3, S5) and abrogates RF2 function (Fig. 4c). On the other hand, the cascade of transitions through the ensemble of intermediate RF2 conformations is efficient for wt-nsECs, and even more so for wt-pre-nsECs where ArfA is fully accommodated to form the ribosomal binding platform for RF2 (Fig. 2b). Because the cascade is the most efficient when the ribosomal binding platform is already pre-accommodated, the intermediate conformations will be even more transiently sampled and only a single E_FRET_ state will be observed before the transition to the collapsed state, as is the case for TCs (Figs. 2 and 4c). These differences in efficiency of the cascade of transitions may explain why, under steady-state X-ray crystallographic and cryogenic electron microscopy (cryo-EM) conditions, ribosome-bound RF2 has been observed in both collapsed and docked conformations in nsECs^27^, but only in the final docked conformation in TCs^6,19,20,23–25,32^. Only time-resolved cryo-EM has captured TC-bound RF2 in the collapsed conformation.^22^

An ensemble of binding intermediates can also be seen in the RF2 dissociation pathway (Figs 2b, 3b, and S2), with the same trend across complexes observed during association replicated here. This suggests that the reverse of the association cascade also acts in the pathway for RF2 dissociation. Interestingly, the rate of RF2 dissociation from the TC we observe here is significantly smaller than that reported in a previous study^21^, leading us to favor a mechanistic model where the translational GTPase class II RF, RF3, is necessary for efficient dissociation of RF2 from TCs. Our model allows us to further propose a mechanism for how RF3 can facilitate this dissociation by engineering a reversal of the cascade through a selective stabilization of the binding intermediates. Corroboration for this model comes from an entirely separate RF2-associated translation pathway, that of post-peptidyl transfer quality control. In this pathway, RF2 and RF3 stably bind to an EC containing a mis-incorporated tRNA and release the aberrant peptide.^43^ It is unknown how this is achieved, since RF2 cannot stably bind to an EC under normal conditions. However, if RF3 stabilizes the binding intermediates that we observed in this study, that would explain how RF3 can both enable RF2 to stably bind an EC with a weak inherent affinity for RF2, and also facilitate RF2 dissociation from a TC where the more stable conformations in the absence of RF3 are the tightly bound collapsed and docked conformations.

It is unknown whether RF3 plays a role in the ArfA-RF2 pathway. Our experiments show that RF2 can dissociate from nsECs at a slightly faster rate than from TCs, leading us to predict that while RF3 might not be essential for this pathway, it may still play a role in the recycling of RF2 from nsECs. Further work is necessary to parse out the exact mechanistic details of RF3 function and how it interacts with the intricate conformational dynamics of RF2 that we observe in TCs and nsECs in our experiments.

Altogether, the work we have presented here highlights the remarkable plasticity of RF2 in catalyzing peptide release from the ribosomal complex in different contexts, an essential step for the proper termination of translation in both canonical and rescue pathways. Equally remarkable is the fact that RF2 undergoes conformational transitions in nsECs that are similar to the ones it undergoes in TCs. This finding highlights the ability of ArfA to functionally replicate interactions that RF2 makes with the TC. The intermediate conformations we observe in this work are relatively short-lived, and this perhaps explains why previous structural studies, which are predicted to underestimate conformational heterogeneity,^44^ did not capture them. Indeed, some of the conformations we report may prove to be too short-lived to be accessed by structural studies and instead may require characterization through kinetic investigations entirely.

Understanding these dynamics and the important conformations these RFs adopt during catalytic activity is critical not only from a mechanistic standpoint but also from one focused on human health as well. Recently, trans-translation has emerged as a potential antibiotic target, with the development of drug leads that can inhibit the process.^18^ For many bacteria, a suitable antibiotic cocktail must target both trans-translation and the ArfA-RF2 pathway in order to be effective. Our work elucidates a potential mechanism by which the ArfA-RF2 pathway may be disrupted. The model of RF2 dynamics that we elucidate here thus forms not only an important part in our understanding of the function of RF2 specifically, and the wider family of class I RFs in general but also highlights a future target for the next generation of broad-spectrum antibiotics.

## MATERIALS AND METHODS

### Construction and purification of Cy5-labeled RF2 constructs

The K12-derived *E. coli* C600 RF2 gene (*prfB*) was cloned into a pProEx-Htb plasmid according to the method previously described for RF1.^45^ To enable overexpression of RF2, the autoregulatory stop codon in *prfB* was removed by deletion of the thymine at nucleotide position 76, leading to the direct translation of aspartic acid at residue position 26 in RF2. Residue 246 was mutated from threonine to alanine to mimic the activity of a B-strain *E. coli*-derived RF2, which is necessary for peptide release activity in MRE600-derived ribosomes.^46^ Labeling mutants of RF2 were engineered in which the native cysteine at position 274 was mutated to serine and the serine at position 184 or the valine at position 271 was mutated to cysteine for the RF2^core^ and RF2^dIII^ constructs, respectively. For both constructs, mutation of the cysteine at position 128 led to protein aggregation, and thus this cysteine was left in its native state given that it has extremely low conjugation efficiency (∼3%).^47^ A pET vector containing the methyltransferase enzyme gene (*prmc*) that methylates RF2 Q252^46,48^ was co-overexpressed with RF2 in BL21-DE3 cells. The 6xHis-tagged RF2 constructs were purified over Ni^2+^-nitrilotriacetic (Ni^2+^-NTA) agarose resin followed by subsequent removal of the 6xHis-tag with Tobacco Etch Virus (TEV) protease cleavage. RF2 activity was verified using a radioactive dipeptide release assay (see below). The Cy5-conjugation reaction was performed by combining a 10-fold molar excess of Cy5-maleimide fluorophore at room temperature for 1 hour in Labeling Buffer [100 mM Tris-acetate (Tris-OAc) pH_RT_ = 7.0, 50 mM potassium chloride (KCl), and 1 mM tris(2-carboxyethyl)phosphine (TCEP)]. The labeled (Cy5)RF2 constructs were further purified from excess fluorophore and unlabeled RF2 using size exclusion chromatography with a HiLoad Superdex 75 column pre-equilibrated in 2x Translation Factor Buffer [20 mM Tris-chloride (Tris-Cl), pH_4°C_ = 7.5, 100 mM KCl, and 10 mM β-mercaptoethanol (βME)] followed by hydrophobic interaction chromatography (HIC) with a TSKgel Phenyl-5PW column (Tosoh Bioscience) using RF2 HIC Buffer A [100 mM sodium hydrogen phosphate/sodium dihydrogen phosphate (Na_2_HPO_4_/NaH_2_PO_4_), pH_RT_ = 7.0, 1 M ammonium sulphate ((NH_4_)_2_SO_4_)] and an increasing linear gradient of RF2 HIC Buffer B (100 mM Na_2_HPO_4_/NaH_2_PO_4_, pH_RT_ = 7.0] over 20 column volumes.

### Construction and purification of ArfA constructs

The gene for ArfA and ArfA^A18T^ (lacking the 17 C-terminal residues to mimic the stem-loop cleavage by RNase III that occurs *in vivo*^49^) was cloned into a pProEx-Htb plasmid and expressed in BL21-DE3 cells. The 6xHis-tagged ArfA constructs were purified over Ni^2+^-NTA agarose resin followed by subsequent removal of the 6xHis-tag with TEV protease cleavage.

### Labeling and amino-acylation of tRNA constructs

*E.coli* tRNA^fMet^ (MP Biomedicals) was aminoacylated with methionine or [^35^S]-methionine using methionyl tRNA synthetase and then formylated using formyltransferase as previously described.^35^ tRNA^Phe^ was labeled at position 47 via the naturally occurring 3-(3-amino-3-carboxypropyl)-uridine residue using a 20-fold molar excess of Cy3 N-hydroxy succinimidyl (NHS) ester (GE Lifesciences) in tRNA Labeling Buffer [50 mM HEPES, pH_RT_ = 8.0, 0.9 M sodium chloride (NaCl)] with 20% labeling efficiency. The Cy3-labeled tRNA^Phe^ was purified from the unlabeled fraction using HIC with a TSKgel Phenyl-5PW column (Tosoh Bioscience) using tRNA HIC Buffer A [10 mM ammonium acetate (NH_4_OAc), pH_RT_ = 6.3, and 1.7 M (NH_4_)_2_SO_4_] and an increasing linear gradient of tRNA HIC Buffer B [10 mM NH_4_OAc, pH_RT_ = 6.3, 10% methanol (CH_3_OH)] as previously described.^35^ Labeled Cy3-tRNA^Phe^ was aminoacylated using phenyl-tRNA synthetase as previously described.^35^

### Preparation of mRNAs

The mRNAs used in this study were derived from the T4 bacteriophage gene product 32 and were in vitro transcribed using T7 RNA polymerase from linearized double-stranded DNA templates, as previously described.^35^ The general mRNA sequence used is 5’-[GG]<u>CAACCUAAAACUUACACA</u>GGGCCC**<u>UAAGGA</u>**AAUAAAA*<u>AUG</u>***(XYZ)_n_**-3’, where the bracketed nucleotides are those necessary for in vitro transcription, the underlined nucleotides are for oligonucleotide hybridization (see below), the bold-underlined nucleotides are the Shine-Dalgarno sequence necessary for ribosome binding, and the italicized-underlined nucleotides represent the start codon AUG (the first codon in the wild-type sequence of T4gp32). The bolded nucleotides in parenthesis referred to as **(XYZ)_n_** refer to the number of codons post-methionine (i.e., codons 2-n) from the wild-type sequence of T4gp32 that are included in the construct, where n=20 for our experiments except for the truncated nsEC constructs. For TCs, codon 3 was mutated to UAA. The differences between the three mRNA constructs are shown below, starting from AUG (codon 1 in wild-type T4gp32):

EC: AUG-UUU-AAA-(4-20)

TC: AUG-UUU-UAA-(4-20)

nsEC: AUG-UUU

For surface immobilization in the smFRET studies, the mRNAs were hybridized to a complementary 3’-biotinylated DNA oligonucleotide (5’-TGTGTAAGTTTTAGGTTGATTTG-Biotin-3’, Integrated DNA Technologies) prior to complex assembly, as previously described.^35^

### Preparation of ribosomal complexes

Ribosomes and translation factors were purified as described previously.^35^ TCs, ECs, and nsECs were prepared in Polymix Buffer [50 mM Tris-OAc, pH_RT_ =7.5, 100 mM KCl, 5 mM magnesium acetate (Mg(OAc)_2_), 5 mM NH_4_OAc, 0.5 mM calcium acetate (Ca(OAc)_2_), 6 mM βME, 5 mM putrescine dihydrochloride, and 1 mM spermidine) as previously described.^35,45^ In all cases, initiation complexes were enzymatically prepared by incubating 1.8 μM ribosome with 2.3 μM initiation factors (IF1, IF2, IF3), 2.7 μM fMet-tRNA^fMet^ (or radioactive [^35^S]-methionyl-tRNA^fMet^ for biochemical studies), 5.4 μM mRNA in presence of 2 mM GTP at 37°C for 20 minutes. The reaction mixture was then combined with an equal volume of 15 μM EF-Tu, 5.4 μM Phe-(Cy3)tRNA^Phe^ or Phe-tRNA^Phe^, 6 μM EF-G, and incubated at 37°C for 10 minutes in order to promote the formation of ribosomal complexes containing fMet-Phe dipeptides that were stalled at the codon in the third position^43^. These ribosomal complexes were then further purified using sucrose density gradient ultracentrifugation (10-40% sucrose w/v) for 12 hours at 25,000 RPM using an SW41 rotor in a Beckman Optima L70 Ultracentrifuge (Beckman) in Buffer C (50 mM Tris-HCl, pH_RT_ = 7.5, 70 mM ammonium chloride (NH_4_Cl), 30 mM KCl, 7 mM magnesium chloride (MgCl_2_), 10 mM βME, 1% beta-D-glucose) for TIRFM experiments, or were buffer exchanged into Buffer C using P6 desalting spin columns (Bio-Rad) for biochemical assays. Purified ribosomal complexes were flash frozen in liquid nitrogen, and stored at −80°C.

### Radioactive dipeptide release assays

Enzymatically assembled ribosomal complexes carrying a [^35^S]-methionyl-Phe-tRNA^Phe^ in the P site were pre-incubated in Buffer C at room temperature for five minutes. Additionally, a mixture of RF2 and 1 mM GTP was separately pre-incubated in the same buffer as the ribosomal complexes at the specified temperature for five minutes. Upon initiation of a time course reaction, the ribosome-containing and RF2-containing mixtures were combined to final concentrations of 100 nM ribosomal complexes, 1 mM GTP, and varying concentrations of RF2. Small samples were taken out at specific time points and quenched with an equal volume of ice-cold 25% formic acid. The desired dipeptides were separated from other products using electrophoretic thin layer chromatography (eTLC) with pyridine-acetate (0.05% pyridine, 20% acetic acid) at 1200 V for 30 minutes^43,50,51^ (Fig. S6a,b). The amount of dipeptide was quantified (Fig. S6c,d,e) using ImageQuant v5.2.

### smFRET experimental and imaging conditions

Microfluidic flow cells were prepared using quartz microscope slides (G. Finkenbeiner) and borosilicate coverslips (VWR) passivated with a mixture of polyethylene glycol and dilute biotinylated polyethylene glycol, as previously described.^50,52,53^ The slide surfaces were functionalized with streptavidin and ribosomal complexes were surface-immobilized via a pre-annealed, biotinylated DNA oligonucleotide tether, as previously reported.^45,50^ The data were collected with a wide-field, prism-based total internal reflection fluorescence (TIRF) microscope with a 100 mW diode-pumped solid-state 532 nm laser (Laser Quantum gem 532) operating at a power of 30 mW (or 15 mW for pre-steady state experiments), measured at the prism, for Cy3 excitation.^45,50,52^ The fluorescence emissions were collected through a 60x, 1.2 NA, water-immersion objective (Nikon), and imaged through a Dual-View (Photometrics) multi-color imaging device using an iXon Ultra 888 electron-multiplying charge-coupled device (EMCCD) camera (Andor) with 2x2-pixel binning and a 100 ms acquisition time per frame. All steady-state experiments were performed at room temperature in Buffer C containing an oxygen-scavenging system composed of 50 U/ml glucose oxidase (Sigma) and 370 U/ml catalase (Sigma), a triplet-state quencher system composed of 1 mM cyclooctatetraene (Sigma) and 1 mM p-nitrobenzyl alcohol (Fluka), and a non-specific interaction blocking system composed of 1 µM duplex DNA oligo and 1 µM BSA.^52^ Pre-steady-state experiments were performed by flowing in (Cy5)RF2^core^ or (Cy5)RF2^core^ and ArfA (1μM) to tethered TCs or nsECs, respectively, at (Cy5)RF2^core^ concentrations of 2.5 nM, 5 nM, 10 nM and 20 nM (prepared by serial dilution from the highest concentration).

### E_FRET_ *vs.* time trajectory analysis

As previously described,^54^ corresponding fields-of-views comprising the fluorescence emissions of hundreds of Cy3 and Cy5 single emitters were aligned using an Iterative Closest Point (ICP) algorithm^55^ to identify the best affine transformation between the two channels. The time-dependent fluorescence emissions from the Cy3 and Cy5 single emitters thus colocalized were fit to Gaussian distributions and extracted to yield fluorescence intensity vs time trajectories. Bleed-through from the Cy3 channel to the Cy5 channel was corrected using a bleed-through parameter (3.5% for all nsECs with (Cy5)RF2^core^, 5% for TCs with (Cy5)RF2^core^, 4% for all complexes with (Cy5)RF2^dIII^). Only trajectories exhibiting anti-correlated Cy3 and Cy5 intensities (*I_Cy3_* and *I_Cy5_*, respectively) and photobleaching (*i.e.* complete loss of fluorescence emission) in a single step were selected and truncated at the time point of Cy3 photobleaching. The corresponding E_FRET_ *vs.* time trajectories were obtained using the equation E_FRET_ = (*I_Cy5_*/(*I_Cy5_* + *I_Cy3_*)^56^ and were used for further analysis.

### Kinetic analysis of pre-steady state E_FRET_ *vs.* time trajectories

Trajectories were analyzed to identify the first binding event of (Cy5)RF2^core^ (defined as the first appearance of *I_Cy5_* and corresponding anticorrelated drop in *I_Cy3_* upon addition of (Cy5)RF2^core^), which were then used to obtain the time to (Cy5)RF2^core^ binding for that individual trajectory. Survival plots of the distributions of binding time so obtained at each concentration of (Cy5)RF2^core^ were fit to exponential decays using Python scripts, using the function *y* = *Ae*^−*kt*^ for the single exponential decay, where *y* is the survival probability at time *t*, *k* is the rate constant, *A* is the normalization constant, and *y* = *Ae*^−*k*_1_*t*^ + *Be*^−*k*_2_*t*^ for the double exponential decay, where *y* is the survival probability at time *t*, *k*_1_ and *k*_2_ are the rate constants for the two phases, *A* and *B* are the corresponding populations for those phases. These yielded the apparent pseudo-first-order rates of association (*k*_*a*_^*app*^s) at these concentrations of (Cy5)RF2^core^. Survival plot fits which yielded an R^2^ value of 0.95 or above for a single exponential fit were taken to be well explained by single exponential decays. The rest were fit using a double exponential decay function. For TCs, the survival plots for RF2 concentrations at 2.5 and 5 nM could be well explained by single exponential decays, while for 10 and 20 nM, the survival plots required at least double exponential decays. The faster phases of the double exponential decays, in these cases, showed a dependence on (Cy5)RF2^core^ concentration and were taken to be the *k*_*a*_^*app*^s at these concentrations. The slower phases, which accounted for a population of ∼10% of the total number of binding times, were found to be orders of magnitude slower than the *k*_*a*_^*app*^s and were likely the result of non-specific binding of (Cy5)RF2^core^ at these higher concentrations. For nsECs, the survival plots for all concentrations could be well explained by single exponential decays. Subsequently, these pseudo-first order *k*_*a*_^*app*^s were plotted as a function of (Cy5)RF2^core^ concentrations, and fit to a linear equation (*y* = *mx* + *c*) to obtain the second order rate of association, *k*_*a*_, for these complexes (Fig. S2).

### Global HMM analysis of steady-state E_FRET_ *vs.* time trajectories

The trajectories for all steady-state experiments for each construct were analyzed using a global variational Bayes HMM analysis, as previously described.^36^ Briefly, the individual E_FRET_ *vs.* time trajectories in the datasets for each ribosomal complex were assumed to be independent and identically distributed (i.i.d.) according to a single global HMM for that complex, which was then inferred using the vbFRET algorithm.^57^ For (Cy5)RF2^core^ datasets and (Cy5)RF2^dIII^ with the Api137-bound TC, a global 2-state HMM was used. For the other (Cy5)RF2^dIII^ datasets, a global 3-state HMM was used for all datasets except for mut-nsEC. In the mut-nsEC, the global 3-state model yielded two zero E_FRET_ states and a very broad non-zero E_FRET_ state encompassing both the extended and the collapsed states. This is likely due to low occupancy of the ribosome-bound non-zero E_FRET_ states of (Cy5)RF2^dIII^ on mut-nsECs, which led to the HMM picking up variations in the zero E_FRET_ state as a separate state instead. To get around this, a 4-state HMM was used, which captured two zero E_FRET_ states, the mid-E_FRET_ state, and the high E_FRET_ state. This 4-state model was collapsed into 3 states by combining both ‘zero’ E_FRET_ states into a single state, *i.e.*, combining the transitions to and from the separate zero E_FRET_ states into that for a single zero state, and the transitions between them into self-transitions of the zero state. These transition probabilities from the HMM transition matrices were used to calculate the rate between specific states using the equation

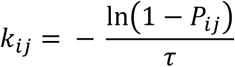

where *k*_*ij*_ is the transition rate between states *i* and *j*, *P*_*ij*_ is the corresponding transition probability, and *τ* is the duration of a single frame.^58^

### Accounting for photobleaching using Api137

The zero E_FRET_ state corresponds to both a ribosome lacking a bound (Cy5)RF2 and one where the Cy5 dye had photobleached; therefore, the rate of transition into the zero state, as calculated by the HMM-based analysis described above, is a sum of the rate of RF2 dissociation and that of Cy5 photobleaching. However, since Api137 can trap RFs in the extended state on the ribosome for extended periods of time that greatly exceed our time window^38,39^, all instances of a drop in E_FRET_ to zero can be attributed solely to photobleaching for an Api137-bound complex. Thus, the rate of transition to the zero E_FRET_ state for this construct was used to estimate the photobleaching rate for all other constructs in our experimental conditions. It should, however, be noted that the rate of photobleaching for any fluorophore is influenced by several factors that can be directly related to the individual complexes themselves. For example, a higher E_FRET_ state for a donor-acceptor pair on a given complex typically corresponds to more (indirect) excitations of the acceptor fluorophore compared with a lower E_FRET_ state for the same pair on a different complex observed under the same experimental conditions. Thus, complexes exhibiting a higher E_FRET_ will likely have a higher rate of photobleaching than those with a lower E_FRET_. Since the Api37-bound TC mostly exhibits the mid E_FRET_ state (∼0.39) of (Cy5)RF2^dIII^, this correction represents a lower bound for the global photobleaching rates, and thus the apparent dissociation rates (k_d_s) calculated using this correction represent a higher bound for the real dissociation rate. It is thus reasonable to expect, for example, that the high E_FRET_ state (∼0.9) for the (Cy5)RF2^core^-bound complexes (or indeed the E_FRET_ states for the closed conformation of (Cy5)RF2^dIII^ (>0.7)) would consequently have a higher rate of photobleaching, and thus the rates of dissociation from these states reported in this work are likely overestimated.

### Construction of post-synchronized 2D histograms

To visualize transitions from all of the non-zero E_FRET_ states to the zero E_FRET_ state for each dataset, the idealized trajectories from the global HMM analysis (using the Viterbi algorithm^57^) were split into dwells in the non-zero E_FRET_ states and dwells in the zero E_FRET_ state using an E_FRET_ threshold of 0.25. All transitions from an E_FRET_ state above 0.25 to one below were collected for the entire dataset and post-synchronized to time zero, which was then used to construct a 2D E_FRET_ vs time histogram. Only a single dwell in the non-zero E_FRET_ states (i.e., E_FRET_ > 0.25) immediately preceding the transition and a single dwell in the zero E_FRET_ states (i.e., E_FRET_ < 0.25) immediately after the transition were considered. Each time-point (frame) in the histogram (i.e., the data points contained in a column at a given time *t*) was normalized to 1. Similarly, for transitions from the zero E_FRET_ state to all of the non-zero E_FRET_ states, a similar procedure was used, except only transitions from an E_FRET_ state below 0.25 to one above were considered. For the histograms showing transitions from the mid-E_FRET_ state to the zero E_FRET_ state and the reverse, only transitions between the mid-E_FRET_ state and zero E_FRET_ state (and single dwells in these states immediately before and after the transitions) were considered.

### Dwell-time analysis of individual E_FRET_ states

The idealized trajectories from the HMMs for each complex yielded the dwell times for individual states. For a given state, these dwells were used to construct a dwell time distribution and construct a survival plot, similar to the procedure for the pre-steady state kinetics. Using Python scripts, these were fit to exponential decays, using the function *y* = *Ae*^−*kt*^ for the single exponential decay and *y* = *Ae*^−*k*_1_*t*^ + *Be*^−*k*_2_*t*^ for the double exponential decay, with the terms defined as above. A threshold of R^2^ = 0.95 for the single exponential fit determined whether the single exponential fit was sufficient to explain the decay or whether at least a double exponential was required. For parsed dwells, the above analysis was restricted to only include dwells of a given state that originated from and resulted in specific states.

## ACKNOWLEDGEMENTS

This work was supported by funds to R.L.G. from the National Institutes of Health (R01 GM084288 and R01 GM137608) and the National Science Foundation (MCB 0644262). B.Y.H. was supported by funds from the NSF Graduate Fellowship (DGE-11-44155).

## AUTHOR CONTRIBUTIONS

N.M., B.Y.H., K.K.R., C.D.K., and R.L.G. designed the research; B.Y.H. performed all the smFRET experiments and collected the data; B.Y.H., N.M., and K.K.R. analyzed the data; N.M., K.K.R, and R.L.G. wrote the manuscript; all of the authors approved the final manuscript.

## DATA AND MATERIALS AVAILABILITY

The open-source Python code for tMAVEN, which was used for the single-molecule data analysis in this work, is freely available via a Git repository (https://github.com/GonzalezBiophysicsLab/tmaven). The EFRET vs time trajectories will be made available upon request to R.L.G.

## SUPPLEMENTAL MATERIALS

**Figure S1.**
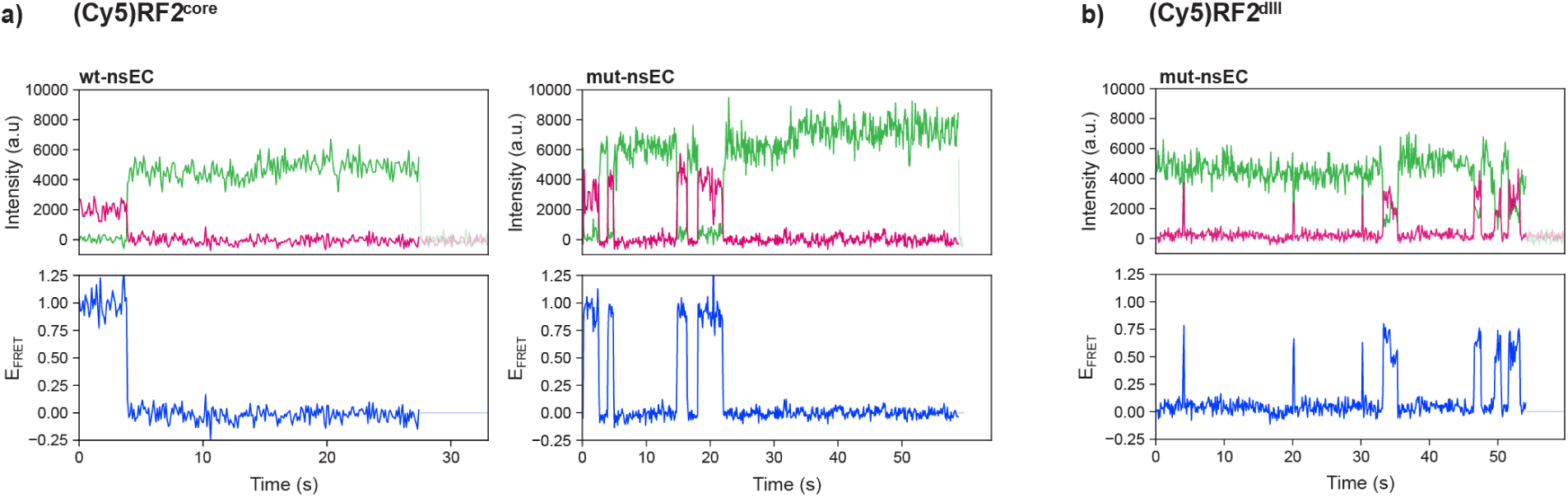
smFRET signals for RF2 stable association and dynamics. Representative smFRET trajectories showing the Cy3 (green) and Cy5 (red) fluorescence intensities and the corresponding E_FRET_ (blue) for **(a)** wt-nsEC (left) and mut-nsEC (right) with the (Cy5)RF2^core^construct and **(b)** mut-nsEC with the (Cy5)RF2^dIII^ construct.

**Figure S2.**
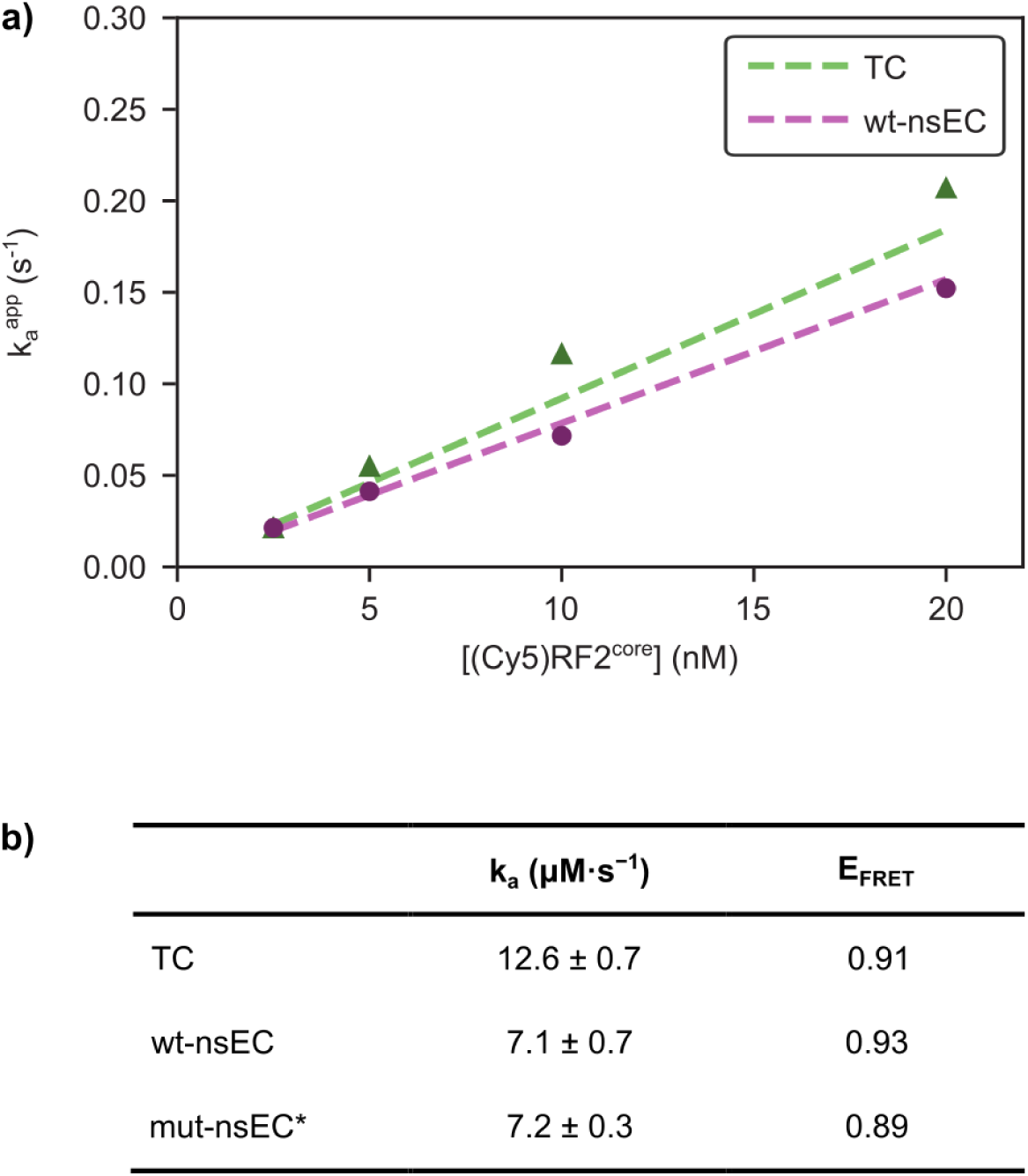
Kinetics of RF2 stable association. **(a)** First-order k_a_ ^app^ for (Cy5)RF2^core^ association with TCs and wt-nsECs, and **(b)** second-order k_a_ for RF2^core^ association with TCs, wt-nsECs and mut-nsECs (*estimated from steady-state experiments).

**Figure S3.**
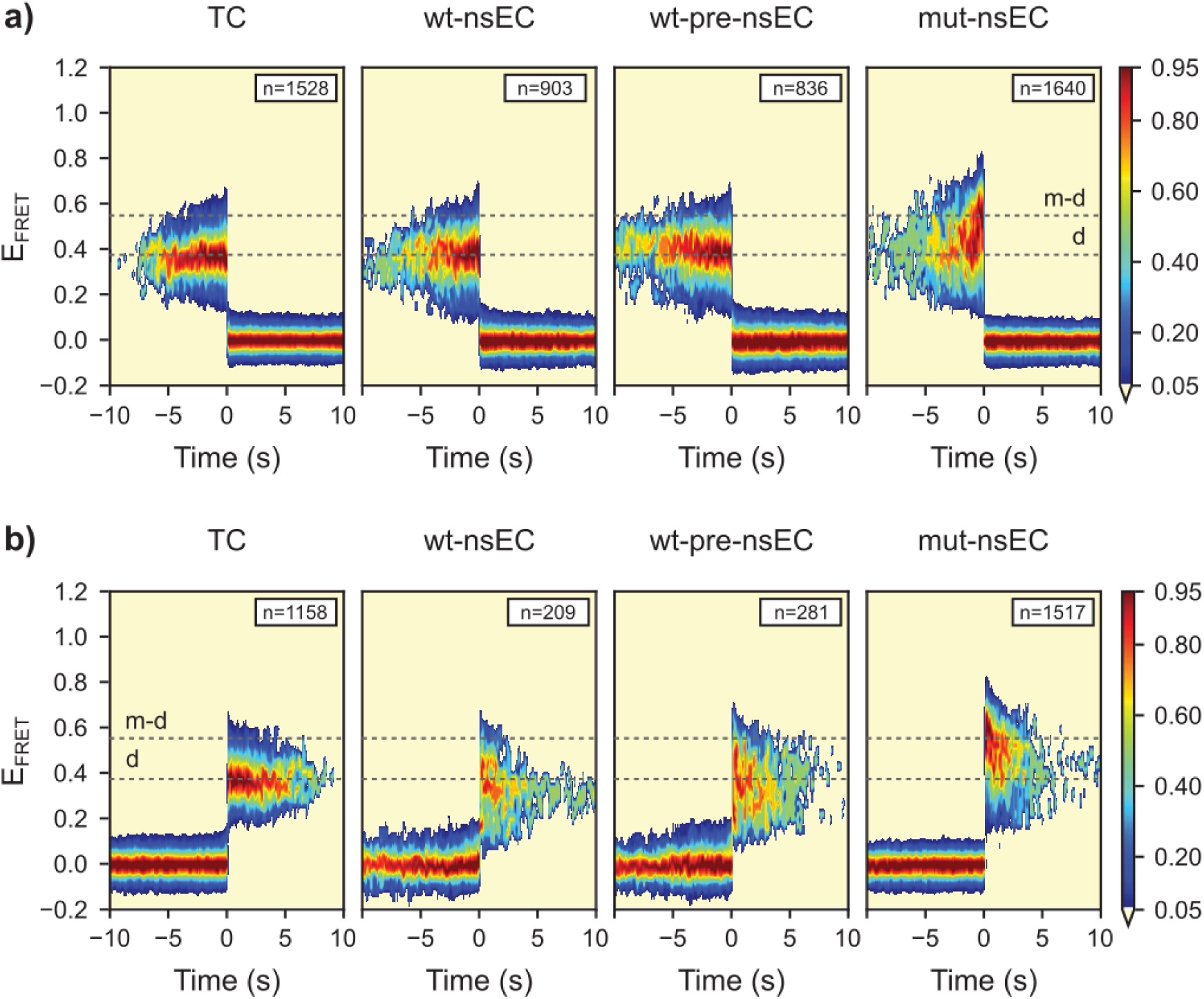
2D E_FRET_ histograms of RF2 dissociation and stable association. 2D histograms post-synchronized to **(a)** the transition from the distal E_FRET_ state to the zero E_FRET_ state, following the dissociation of ribosome-bound (Cy5)RF2^dIII^, and **(b)** the transition from the zero E_FRET_ state to the distal E_FRET_ state, following the stable association of (Cy5)RF2^dIII^ to the ribosome, for different ribosomal complexes. Grey dashed lines show the E_FRET_ states corresponding to the distal (d), and transient mid-distal (m-d) conformations of (Cy5)RF2^dIII^.

**Figure S4.**
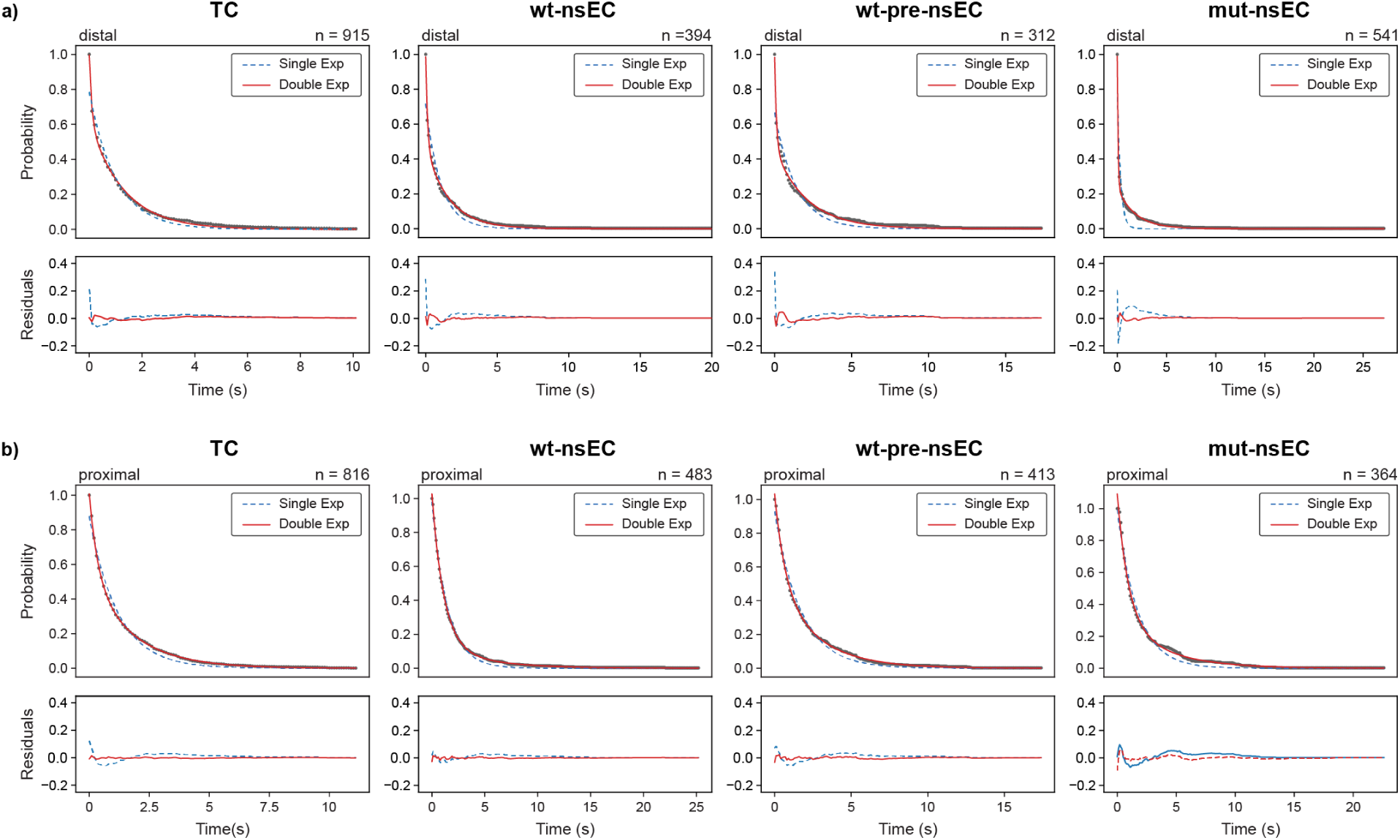
Dwell analysis of the extended and collapsed conformations of ribosome-bound (Cy5)RF2^dIII^. Survival plots of the dwell time distributions (black) along with fits of single-exponential (blue) and double-exponential (red) decays (above) and the corresponding residuals (below) for **(a)** the distal state, and **(b)** the proximal state for different ribosomal complexes.

**Figure S5.**
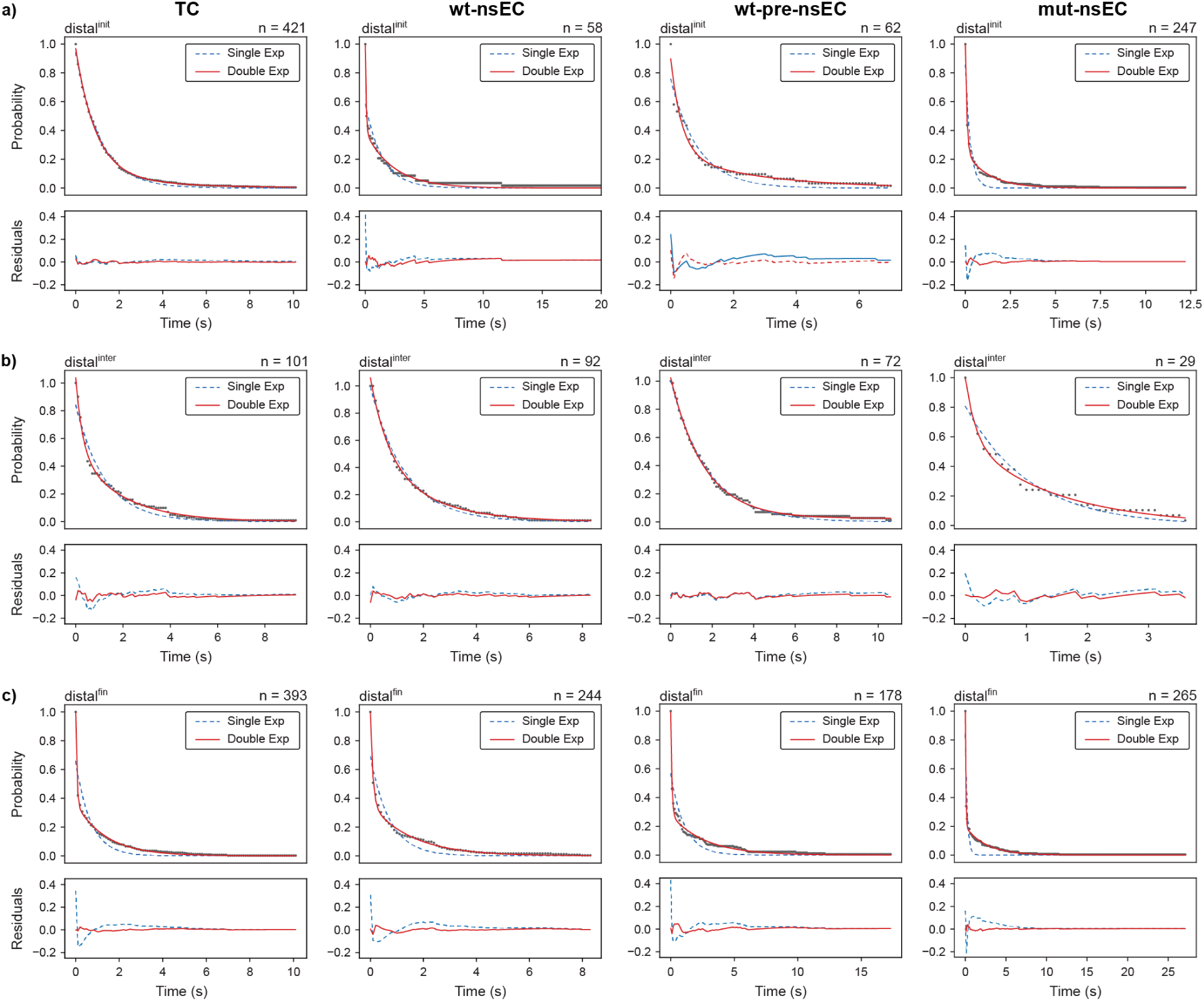
Parsed dwell analysis of the extended conformation of ribosome-bound (Cy5)RF2^dIII^. Survival plots of the dwell time distributions (black) along with fits of single-exponential (blue) and double-exponential (red) decays (above) and the corresponding residuals (below) for **(a)** distal^init^ **(b)** distal^inter^, and **(c)** distal^fin^, for different ribosomal complexes.

**Figure S6.**
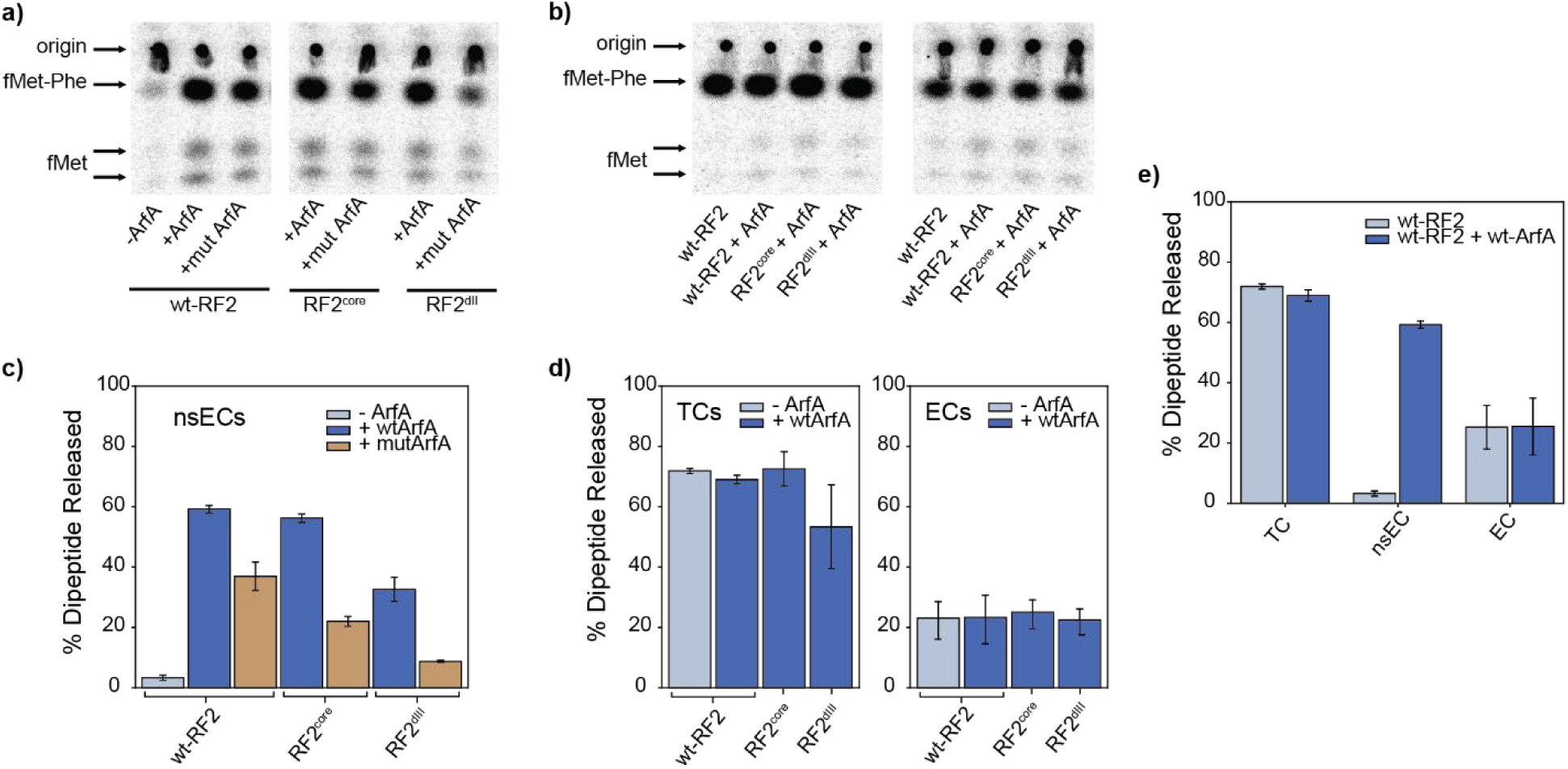
fMet-Phe dipeptide release assay. **(a)** representative eTLC of fMet-Phe dipeptide released from nsEC complexes incubated with wt-RF2, (Cy5)RF2^core^, or (Cy5)RF2^dII^ and either no ArfA (for wt-nsEC only), wt-ArfA, or mut-ArfA after one minute at 37°C; **(b)** representative eTLC of fMet-Phe dipeptide released from TC or EC complexes incubated with wt-RF2, (Cy5)RF2^core^, or (Cy5)RF2^dII^, with (for wt-RF2 only) and without wt-ArfA, for one minute at 37°C; **(c)** % dipeptide released from nsECs, quantified from eTLC analyses performed as in (a); **(d)** % dipeptide released from ECs and TCs, quantified from eTLC analyses performed as in (b); **(e)** comparison of % dipeptide released from TCs, nsECs, and ECs with wt-RF2, with or without wt-ArfA, from eTLC analyses performed as in a) and b).

**Table S1.** E_FRET_s observed for all complexes. E_FRET_s obtained from our HMM-based analysis for complexes with (Cy5)RF2^core^ and complexes with (Cy5)RF2^dIII^. Errors represent the standard error of the mean, as derived from our HMM analysis.

|  | (Cy5)RF2<sup>core</sup> | (Cy5)RF2<sup>dIII</sup> proximal | (Cy5)RF2<sup>dIII</sup> distal |
| --- | --- | --- | --- |
| **TC** | 0.91 ± 0.009 | 0.76 ± 0.009 | 0.37 ± 0.006 |
| **wt-nsEC** | 0.93 ± 0.015 | 0.70 ± 0.009 | 0.36 ± 0.007 |
| **wt-pre-nsEC** | 0.91 ± 0.015 | 0.69 ± 0.009 | 0.37 ± 0.007 |
| **mut-nsEC** | 0.89 ± 0.009 | 0.75 ± 0.009 | 0.43 ± 0.009 |
| **Api137-bound TC** | - | - | 0.39 ± 0.004 |

**Table S2.** Rates obtained from parsed dwell analysis of the extended conformation of ribosome-bound (Cy5)RF2^dIII^. Rates, amplitudes, and R^2^ values obtained from our parsed dwell analysis. Note that for wt-nsEC distal^init^, the R^2^ for the double exponential fit is 0.94, which is lower than our threshold value of 0.95. However, the single exponential fit was significantly worse, with an R^2^ value of 0.84; thus, a double exponential fit was assigned in this case.

| | | $k_1$ (s<sup>-1</sup>) | $k_2$ (s<sup>-1</sup>) | $A_1$ | $A_2$ | $R^2$ |
| --- | --- | --- | --- | --- | --- | --- |
| **TC** | distal<sup>init</sup> | 0.09 | - | 0.94 | - | 0.99 |
|  | distal<sup>inter</sup> | 0.08 | - | 0.84 | - | 0.96 |
|  | distal<sup>fin</sup> | 0.07 | 1.97 | 0.36 | 0.63 | 0.99 |
| **wt-nsEC** | distal<sup>init</sup> | 0.04 | 1.36 | 0.39 | 0.60 | 0.94* |
|  | distal<sup>inter</sup> | 0.07 | - | 0.99 | - | 0.99 |
|  | distal<sup>fin</sup> | 0.06 | 1.10 | 0.36 | 0.63 | 0.99 |
| **wt-pre-nsEC** | distal<sup>init</sup> | 0.04 | 0.28 | 0.24 | 0.66 | 0.97 |
|  | distal<sup>inter</sup> | 0.06 | - | 1.00 | - | 0.99 |
|  | distal<sup>fin</sup> | 0.04 | 1.13 | 0.28 | 0.71 | 0.98 |
| **mut-nsEC** | distal<sup>init</sup> | 0.08 | 1.18 | 0.25 | 0.74 | 0.99 |
|  | distal<sup>inter</sup> | 0.07 | 0.59 | 0.57 | 0.42 | 0.99 |
|  | distal<sup>fin</sup> | 0.04 | 1.55 | 0.20 | 0.80 | 0.99 |

\* highest $R^2$ value obtained for this complex

